# The Effects of Prelimbic Inhibition on Time-based Intervention and Impulsive Choice

**DOI:** 10.64898/2026.01.08.698432

**Authors:** Maxwell J. McLeod, Kelsey Panfil, Lexe West, Ian Davis, Cole Vonder Haar, Kimberly Kirkpatrick, Travis R. Smith

## Abstract

Impulsive choice is the suboptimal preference for a smaller-sooner (SS, “impulsive”) option over a larger-later (LL, “self-controlled”) option. Fixed-interval (FI) training delivers delayed-reinforcement trials to increase LL choices and improve FI timing precision. While there are plenty of studies exploring the neurobiological factors underlying impulsive choice, it is unknown what neurobiological changes account for the FI training effects. The prelimbic cortex (PL) region is implicated in both impulsive choice and timing. To investigate the role of the PL, we used designer receptors exclusively activated by designer drugs (GiDREADDs) to reversibly inhibit the PL during either the FI training phase or the follow-up impulsive choice task in male and female Sprague-Dawley rats. Compared to a control group, the GiDREADDs rats showed reduced LL choices when CNO was administered during the FI training or impulsive choice tasks. GiDREADDS did not alter response rates or latency to choose. Overall, these data demonstrate that inhibition of the PL increases impulsive choice and may block the effect of the FI training to improve self-control.

## 1. Introduction

Impulsive choice involves choosing a smaller reinforcer that is available after a short delay (the smaller-sooner, or SS) versus a larger reinforcer that is available after a longer delay (the larger-later, or LL;Mazur, 2000; Odum, 2011). Preference for the SS option is an indicator of impulsive choice and a predictor of maladaptive behaviors and conditions such as attention deficit hyperactivity disorder (ADHD) (e.g., Barkley et al., 2001), substance use disorders (SUDs) (e.g., Amlung et al., 2017), pathological gambling (e.g., Bickel et al., 2012; Reynolds, 2006), and obesity (e.g., Bickel et al., 2021). There is interest in exploring the underlying psychological and neurobiological sources of impulsive choice to develop and refine interventions to reduce impulsivity and potentially alleviate associated conditions.

In humans (Baumann & Odum, 2012) and rats (Marshall et al., 2014) individuals that perceive time intervals more precisely and/or accurately are more likely to make self-controlled choices. The impulsive choice task inherently involves delays; thus, imprecise interval timing may result in suboptimal choice behavior. Deficits in interval timing abilities are also implicated in ADHD (Bluschke et al., 2018) and substance use disorders (Paasche et al., 2019) further establishing a relationship between impulsive choice and timing. Therefore, interventions that improve self-control and interval timing abilities may potentially target an underlying mechanism that may address these maladaptive conditions (Smith et al., 2019). Time-based behavioral interventions provide delay exposure using a fixed-interval (FI) schedule to improve self-control and timing performance (Peterson & Kirkpatrick, 2016; Smith et al., 2015; Smith et al., 2024). FI training is long-lasting, generalizes across choice tasks (Bailey et al., 2018), and can improve self-controlled choices with as few as 400 training trials (Panfil et al., 2024).

Research on the relationship between timing and impulsive choice has drawn attention to brain regions that subserve both functions. Cellular activity responding to interval timing (seconds to minutes range) is present in prelimbic, premotor/sensori-motor, and parietal cortices, hippocampus, and striatum (Tallot & Doyère, 2020). Among those regions, the prelimbic (PL) is involved in both interval timing and impulsive choice. In the context of interval timing, PL pyramidal cells demonstrate ramping firing activity during specific delay periods, which may be a fundamental cortical timing mechanism (Emmons et al., 2017; Kim et al., 2013; Tiganj et al., 2017; Xu et al., 2014). Additionally, inactivation of PL disrupts ramping activity (Emmons et al., 2019; Parker et al., 2015) and decreases timing precision (i.e., wider peak spread) on peak interval tasks (e.g., increases timing error; Buhusi et al., 2018; Parker et al., 2015). PL activity predicts timing errors in temporal bisection tasks (Oshio et al., 2006) and disruption of PL activity also impairs temporal discrimination (Kim & Zauberman, 2009; Nepomoceno et al., 2024; Parker et al., 2015).

The PL is a key prefrontal region involved in processing temporal information and is important in impulsive choice tasks that necessarily involve delays. Sackett et al. (2019) reported that a population of PL neuronal activity showed phasic responses to both LL and SS choices. The balance shifted toward SS-responsive neurons with increasing LL delays and a concomitant increase in SS choices in the free-choice trials was observed. The more impulsive rats in the study sample showed relatively more SS responsive neurons. These correlations suggest that the PL may encode choice value. Wenzel et al. (2023) demonstrated a causal relationship by inhibiting projections from the PL to the nucleus accumbens core and observed a decrease in LL choices. Other studies have disrupted PL functioning and have also reported reductions in proportion of LL choices (e.g., Churchwell et al., 2009; Yates et al., 2014).

The present study aimed to experimentally inhibit the PL during either the FI training phase or the subsequent impulsive choice task using inhibitory (Gi) designer receptors exclusively activated by designer drugs (Gi-DREADDs). Rats underwent surgery to receive infusions of either a Gi-DREADD virus or a control virus expressing GFP, which is inert with respect to neuronal activity with or without a designer drug present. Rats then received the designer drug clozapine N-oxide (CNO) to inhibit DREADD-expressing neurons, administered either during FI training or during the impulsive choice task. If FI training enhances self-control through neuroplastic changes in PL-centered timing circuits, then CNO administration during FI training should reduce large-later (LL) choices compared to rats given CNO without Gi-DREADD expression.

## 2. Methods

### 2.1. Animals

Mixed-sex Sprague Dawley rats (*N* = 72, 36 males, 36 females; Charles River, Stone Ridge, NY) were used in this experiment. Rats were tested in four separate cohorts (*n* = 12 in cohorts 1 and 2; *n* = 24 in cohorts 3 and 4). They arrived at approximately 30 days of age (51-75 g in weight) and received their surgery at approximately 90 days of age. The rats were housed in same-sex pairs and maintained on a 12-hr light:dark schedule (lights off at approximately 7 am). The rats were tested during the dark phase of the cycle. Rats had access to water *ad libitum* in the home cages and in the experimental chambers and were food restricted and maintained at a target of 85% of their projected *ad libitum* weight, as derived from growth-curve charts obtained from the supplier.

### 2.2. Apparatus

The experiment was conducted in 24 operant chambers (Med-Associates, St. Albans, VT), each housed within a sound-attenuating, ventilated box (74 × 38 × 60 cm). Each chamber (25 × 30 × 30 cm) was equipped with a stainless-steel grid floor; two stainless steel walls (front and back); and a transparent polycarbonate side wall, ceiling, and door. Two pellet dispensers, mounted on the outside of the front wall of the operant chamber, delivered 45-mg food pellets (Bio-Serv, Flemington, NJ) to a food cup centered on the lower section of the front wall. Head entries into the food magazine were detected by an infrared photobeam. Two retractable levers were located on opposite sides of the food cup. The chamber was also equipped with a house light centered at the top of the chamber’s front wall, as well as two nose-poke key lights that were each located above the left and right levers. Water was always available from a sipper tube that protruded through the back wall of the chamber. Experimental events were controlled and recorded with 1-ms resolution by the software program MED-PC V.

### 2.3. Procedure

Rats were randomly assigned to four groups in a 2 (virus) × 2 (phase) factorial design prior to experimentation. The virus factor comprised two groups based on the virus infused into the prelimbic cortex: inhibitory (Gi) DREADD or control GFP (labeling). The phase factor comprised two groups defined by whether they received CNO (1 mg/kg, i.p.,10 minutes prior to testing) either during FI Training or Impulsive Choice task. This design controlled any potential endogenous effects of CNO.

Figure 1 shows the experimental timeline. Rats were initially trained to collect pellets from a receptacle and to lever press for pellet delivery. Subsequently, rats underwent surgeries to infuse either the Gi DREADD or GFP virus into the prelimbic cortex and were allowed six weeks for viral expression. Following recovery, rats completed an FI Training phase consisting of nine sessions. Next, all rats completed an abbreviated SS-delay impulsive choice task over nine sessions.

**Figure 1.**
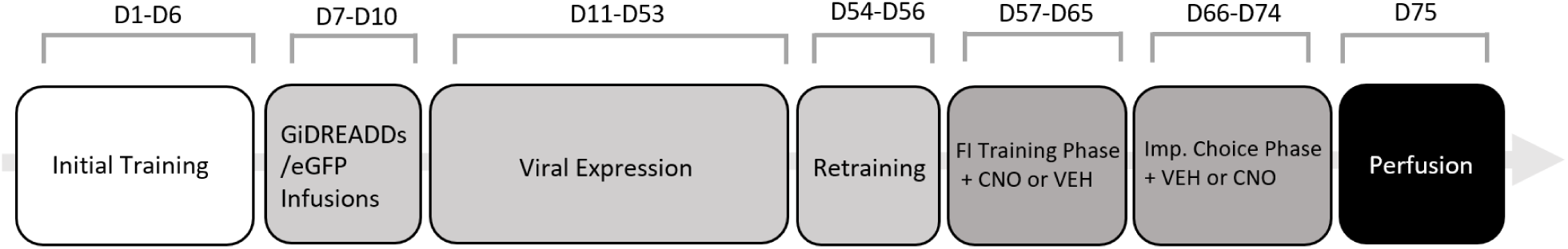
Experimental timeline showing the progression of the study

#### 2.3.1. Surgical Procedure

Rats underwent surgery to transduce an active inhibitory Gi DREADD (AAV8-CaMKlla-hM4D[Gi]-mCherry) or a sham labeling virus (AAV8-CaMKlla-EGFP) into the PL. Rats were anesthetized with isoflurane (3-5% induction, 1-3% maintenance) in 0.5L/min oxygen. Rats received 3-5 ml injection of warm saline (s.c.) 2% lidocaine (0.25 ml/kg s.c.) at the incision site , and meloxicam (2 mg/kg, s.c.). The surgical site was sanitized prior to midline incision. Bilateral burr holes allowed insertion of a microinfusion needle attached to a 1.0 μl syringe infused 0.5 μl (0.1 μl/min with 5 min dwell time before extracting needle) of active Gi DREADD expressing virus or sham virus bilaterally into the PL, +3.2 mm AP, ± 0.8 mm ML, -2.8 mm DV. Rats were maintained six weeks prior to behavioral assessment for recovery and full expression of the inhibitory receptors.

#### 2.3.2. Initial Training

The rats received 4 days of initial training, which consisted of 1 session of magazine training and 3 sessions of lever press training. Magazine training involved the delivery of food pellets to the food cup on a random-time 60-s schedule. Following magazine training, the rats were trained to press both the left and right levers. Initially, the food pellets were delivered on a fixed-ratio (FR) 1 schedule of reinforcement until 20 food pellets were delivered for responding on each lever. The FR 1 was followed by a random ratio (RR) 3 schedule of reinforcement, where 3 responses are required, on average, per reinforcer; which lasted until 20 reinforcers were delivered for responding on each of the two levers. The RR 3 was followed by an RR 5, which lasted until the rats earned 20 food pellets for responding on each of the two levers. Pre-training sessions lasted for up to 2 hr.

#### 2.3.3. General Experimental Procedure

Figure 2 shows the standard behavioral procedures used in FI Training and Impulsive Choice tasks. FI training and impulsive choice trials followed similar behavioral procedures. Trials started with the illumination of the house-light and lever(s) extended into the chamber. The first press (“selection response”) to a given lever extinguished the house-light, retracted the unselected alternative lever (if it was presented), illuminated a cue-light above the selected lever, and started the delay timer. The FI schedule allowed responses during the delay without any consequences. After the programmed delay elapsed, pellet delivery was primed and the first lever press (“collection response”) after the delay resulted in pellet delivery, retraction of the lever, extinguishing of the associated cue-light, and the start of a 60-s intertrial interval (ITI). Responding on the LL lever resulted in 2 pellets after a 30-s delay. Responding on the SS lever resulted in 1 pellet after an *x*-s delay. The SS delay (*x*) varied across conditions (see below). Left/right lever assignment of the SS/LL options was counterbalanced. FI training sessions and forced-choice trials during the impulsive choice task extended one lever into the chamber during the trial. Free-choice trials extended both levers into the chamber. The FI Training consisted solely of forced-choice trials, where only one option was delivered.

**Figure 2.**
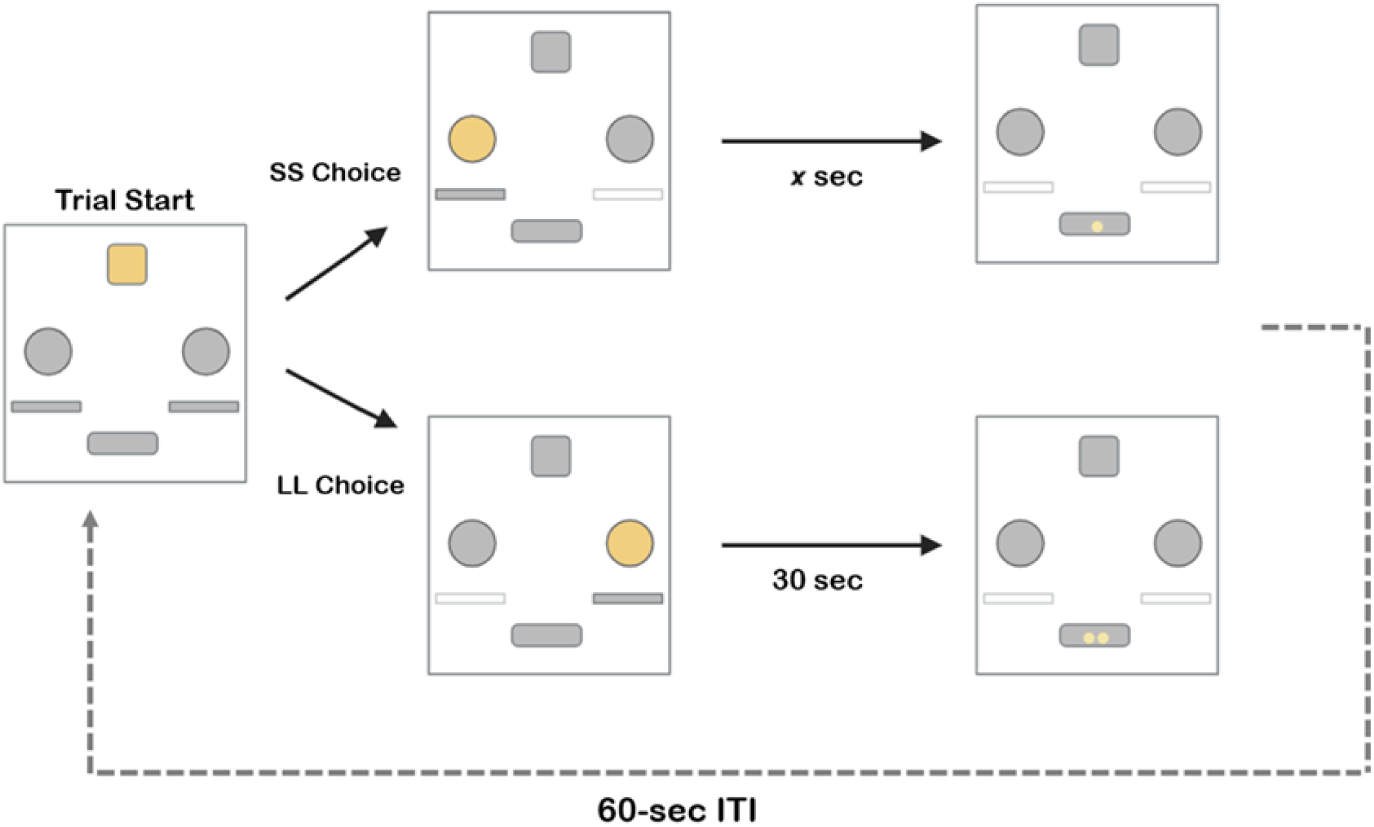
Illustration of the trial structure used in FI Training and Impulsive Choice tasks. Trial started with houselight illumination, choice illuminated a cue-light above the lever, FI responding transitioned to pellet delivery, and a 60-s ITI followed. The SS delay (*x*) varied across conditions.

##### 2.3.3.1. Fixed Interval Training

The FI training trials proceeded as described in Figure 2, except all trials forced exposure to a single lever representing the SS and LL options across blocks of sessions. Sessions lasted until 50 pellets were earned or 70 minutes after CNO/Vehicle dosing. Rats received 9 sessions of FI training exposing them to 6 consecutive sessions on the LL contingencies (2 pellets after a 30-s delay) and 3 consecutive sessions on the SS contingencies (1 pellet after a 10-s delay; i.e.,*x* = 10). FI training sessions on the LL lever included 25 trials to earn 50 pellets and only the LL lever was presented. FI training sessions on SS lever included 50 trials to earn 50 pellets and only the SS lever was presented. By the end of FI training, a total of 300 training trials (150 on SS, 150 on LL) were delivered. Order of exposure to the LL and SS lever during FI training was counterbalanced across rats.

##### 2.3.3.2. Impulsive Choice Procedure

Rats completed the Impulsive Choice task for 9 sessions. The SS and LL trials proceeded as described in the general procedure (Figure 2), and the SS delay (*x*) increased across sessions. Each session started with 6 SS forced-choice trials and proceeded with a block of 36 trials included a random mixture of 24 free-choice and 12 forced-choice trials (6 on LL lever and 6 on SS lever). The first 5 sessions delivered a 10-s SS. The SS delay (*x*) then increased by 5 s for each subsequent session (15, 20, 25, and 30 s). The SS and LL assigned levers were the same as in the FI training. Sessions lasted for 42 trials or 70 minutes after CNO/vehicle dosing.

#### 2.3.4. Perfusion and Virus Placement Assessment

Rats were transcardially perfused used to deliver a 0.9% saline solution to flush out the blood and then delivered a 4% paraformaldehyde solution to fix the tissue. Slices were mounted and the PL imaged for the fluorescent tag for the active Gi DREADD (mCherry) and the GFP labeled control. After losses and exclusions, the final sample consisted of 31 rats (GFP & CNO in Choice: *n* = 6, 3 male; GFP & CNO in FI Training: *n* = 6, 3 male; GiDREADD & CNO in Choice: *n* = 9, 5 male; GiDREADD & CNO in FI Training: *n* = 10, 6 male).

### 2.5. Data Analysis

The primary analysis investigating impulsive choice used a generalized linear mixed effects model (glmer function in R) to fit the proportion of LL choices (specifying a binomial response distribution, 1 = LL choice, 0 = SS choice; i.e., logistic regression) as a function of SS delay (*x*), virus, and phase. Virus and phase a were effect-coded and SS delay (10-30) was scaled between 0 and 1 where 0 represents the 10-s SS delay and 1 represents the 30-s SS delay with the 10-s SS delay serving as the intercept. All interactions were included. The random-effects included individual subjects with a random intercept to account for individual differences. All choices and sessions were included in the analysis, except for the first two sessions when early learning of the choice task was occurring.

A response time analysis measured the latency between lever insertion and the choice response at the start of the trial. A linear effects model (lmer function in R) fitted the log-transform of the response latency (to get a normal distribution) as a function of phase, virus, and their interaction. Only data at the 10-s delay was included because that is where the largest choice differences were observed. Like the Impulsive Choice model, the first two sessions were excluded. The random effects included individual subjects with a random intercept to account for individual differences.

For FI timing analysis, the response rate (responses/sec) data for the LL delay during the choice task was fitted with a cumulative normal distribution function (i.e., sigmoidal function) to model responses across the FI 30-s delay (there were insufficient observations to model the responses during SS delays). A nonlinear multi-level model (nlme function in R) assessed differences in FI response rates as a function of virus and phase. The random effects included individual subjects with a random intercept to account for individual differences. The following equation modeled the data:

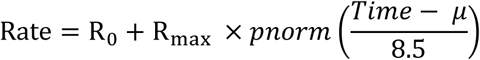

Where Rate is responses per second (dependent measure), Time is the time in trial (independent measure), and *pnorm* is a function for the cumulative normal distribution. Free parameters include R_0_ (baseline response rate), R_max_ (change in response rates from baseline to asymptotic maximum rate), and µ (center of the sigmoidal curve, or inflection point). The 8.5 represents a fixed standard deviation parameter (σ) to avoid multicollinearity that was observed between σ with µ.

## 3. Results

Figure 3 shows the PL Gi DREADD expression from a representative sample (A) and a group-level expression map of viral fluorescence at +3.2 mm anterior (the infusion site) and +4.2 mm anterior from Bregma.

**Figure 3.**
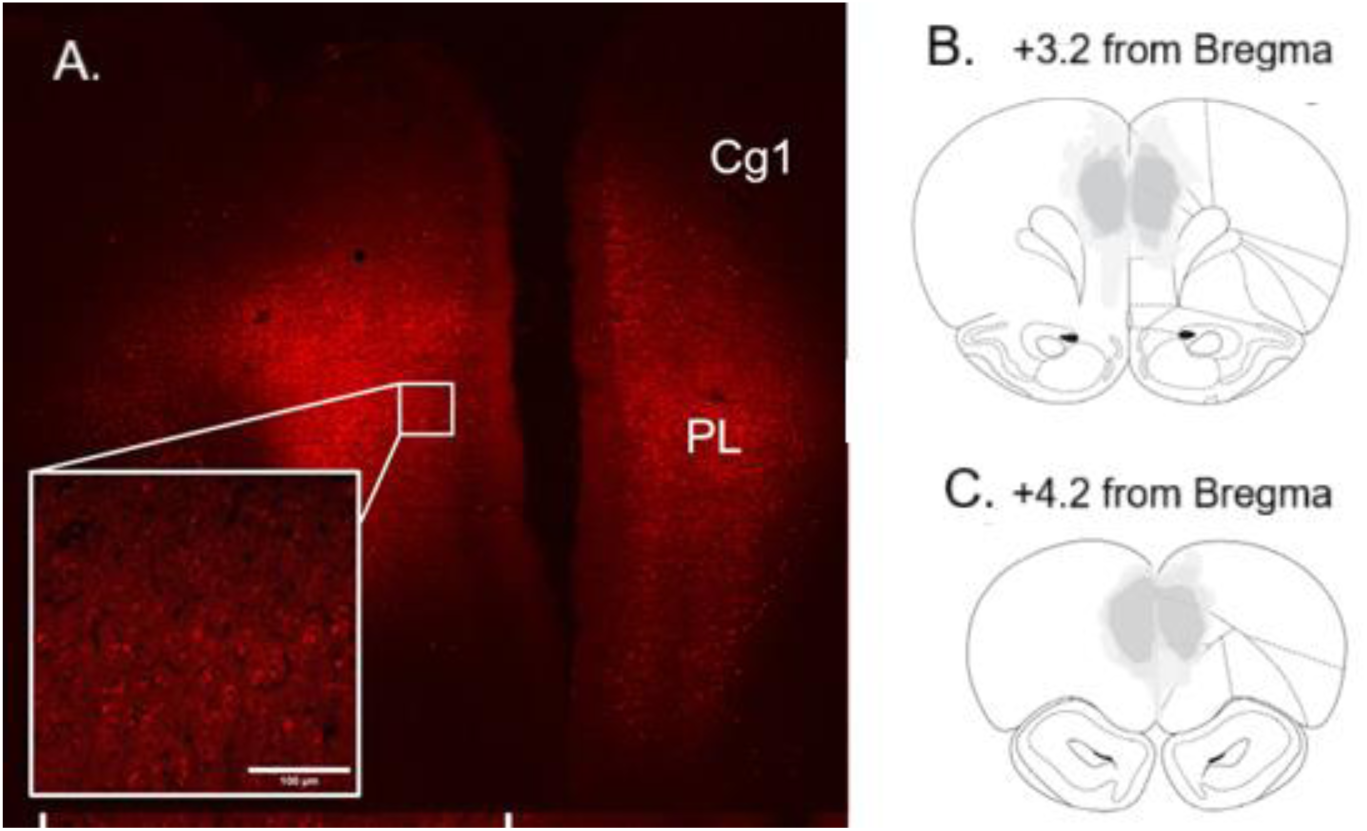
PL DREADDs expression. (A) Example of DREADDs expression from section around +3.2 AP from bregma. (B) Average viral spread of DREADDs overlaid on atlas +3.2 from bregma (infusion site). (C) Expression +4.2 from bregma.

### 3.1. Impulsive Choice

Figure 4 shows the proportion of LL choices (i.e., self-controlled preference) across SS delays for the virus and phase conditions. There was a main effect of SS delay, as delays increased across sessions the proportion of LL choices increased (*b* = 2.96, *SE* = 0.14, *z* = 20.92, *p* < .001). There was a main effect of virus where the inhibitory Gi DREADDs groups made fewer LL choices compared to the GFP virus groups at the 10-s SS delay (*b* = - 0.78, *SE* = 0.33, *z* = -2.38, *p* = .02). There was a three-way interaction between SSDelay, phase, and virus where the differences in slope (SSDelay) between inhibitory DREADD and control virus was different depending upon whether CNO given in the choice or intervention phase (*b* = -0.55, *SE* = 0.14, *z* = -3.90, *p* < .001). A post-hoc test revealed that the sensitivity to the SS delay was greater for the Gi DREADDs group compared to the GFP control when CNO inhibited the PL in the FI Training (*b* = 1.48, *SE* = 0.37, *z* = 4.02, *p* < .001), but not when CNO was delivered during the Impulsive Choice task (*p* = 0.09). Post-hoc tests revealed that the mean proportion of LL choices at the 30-s SS delay was lower for the Gi DREADD group whenCNO was administered in the Impulsive Choice task (*b* = -2.11, *SE* = 0.98, *z* = -2.14, *p* = .03), but not the FI Training task (*p* = 0.80).

**Figure 4.**
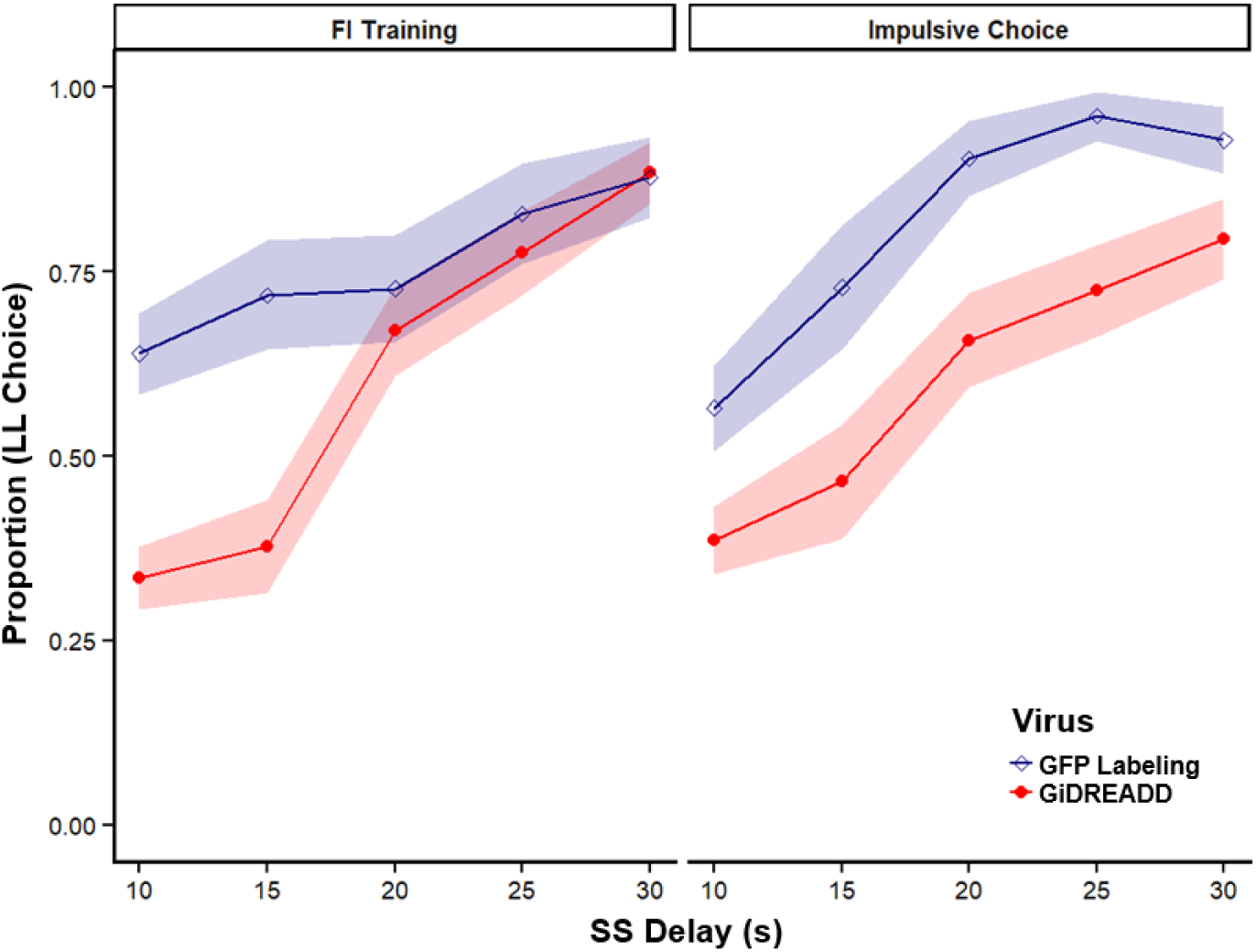
Proportion LL choice across the SS delay between the active Gi DREADD group (red, filled circles) and the GFP Labeling control group (blue, open diamonds). Left panel shows CNO dosing effects during FI Training and the right panel shows CNO effects during the Impulsive Choice procedure. Ribbons represent ±SEM.

### 3.2. Choice Reaction Times

Figure 5 displays the choice reaction times for SS and LL choices during the 10-s SS delay testing. The model did not find any differences between groups (*p*s > .06).

**Figure 5.**
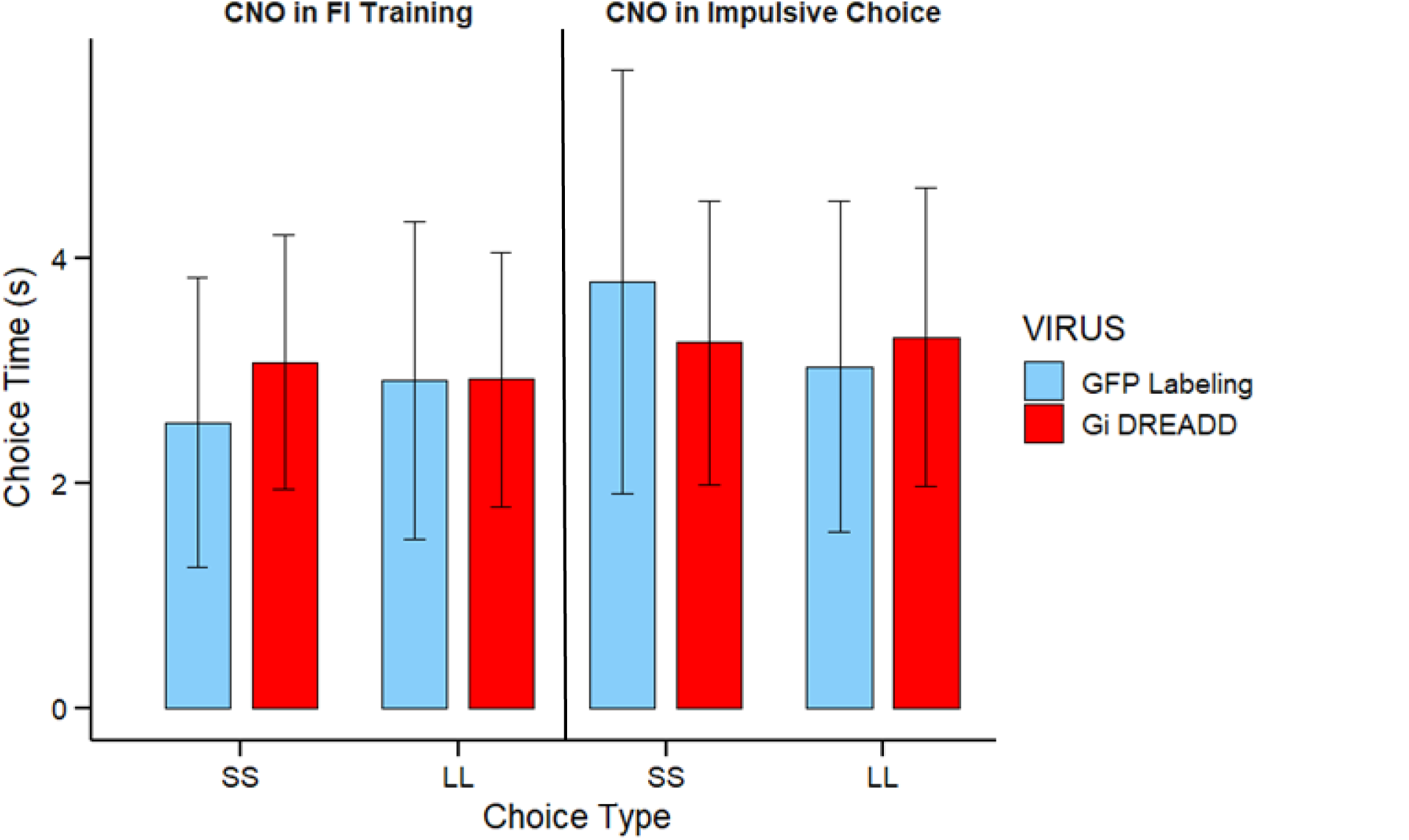
Choice reaction time (s) as a function of choice (SS or LL), Virus groups, and CNO dosing groups.

### 3.3 Choice FI *Response Rates*

The model assessing response rates during the LL delay during the choice task did not find any differences on the R_0_ and R_max_ parameters between the groups (*p*s > .17), as shown in Figure 6. The µ parameter (inflection point in the interval where response rates slow to negative acceleration) showed a virus x phase interaction (*b* = -2.39, *SE* = 0.82, *z* = -2.89, *p* = .004). A post-hoc analysis showed an earlier inflection point for the Gi DREADD group (compared to GFP control) when CNO was administered in the Impulsive Choice task (*b* = -5.22, *SE* = 2.53, *z* = -2.07, *p* = .04), but a later inflection point when CNO was administered in the FI Training task (*b* = 4.34, *SE* = 2.14, *z* = 2.03, *p* = .04).

**Figure 6.**
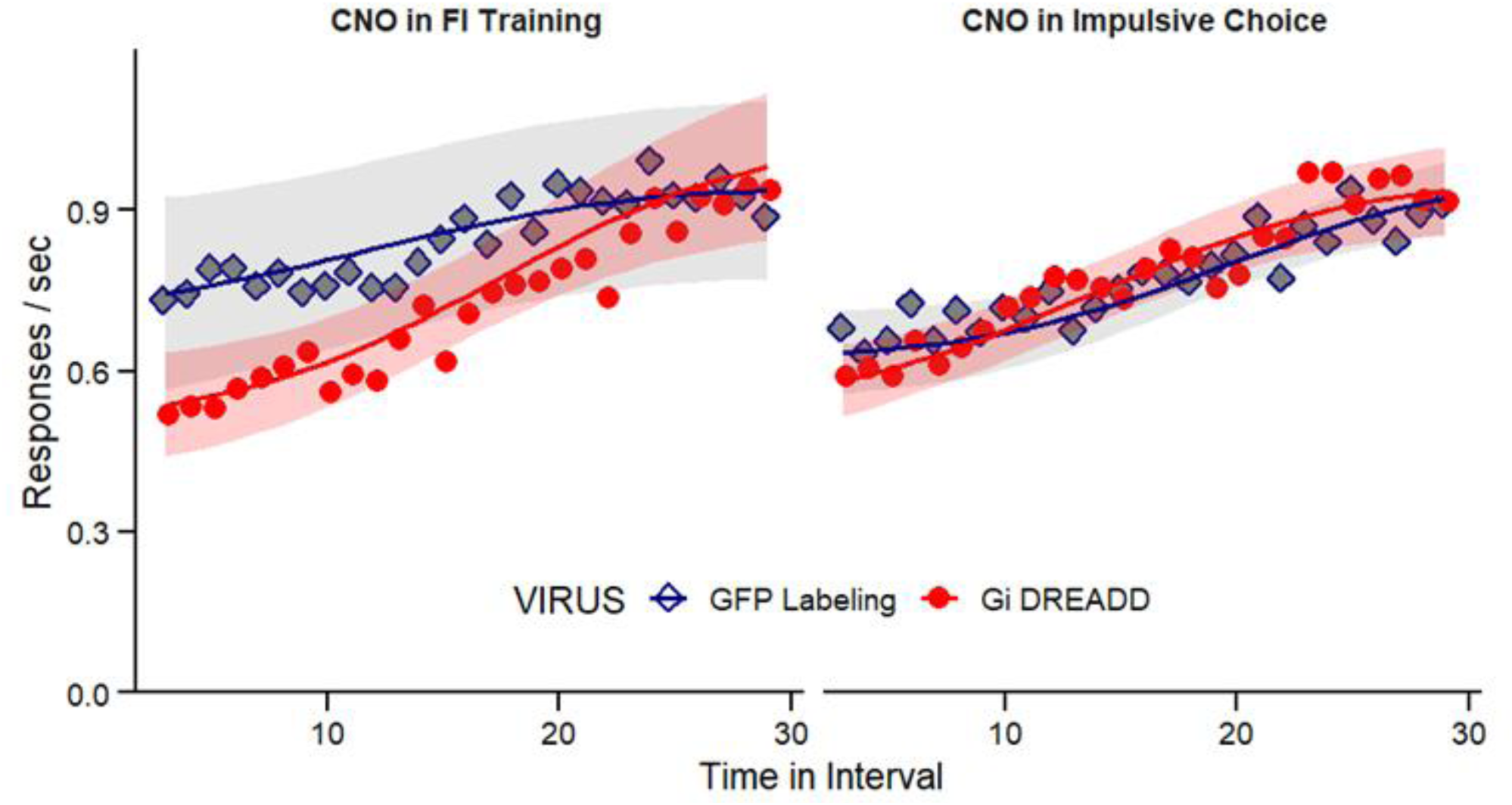
Mean response rate (presses per second) during LL delays in the impulsive choice task as function of time in the interval between Gi DREADD (red circles) and GFP Labeling (blue diamonds) groups. Left panel shows CNO dosing effects during Impulsive Choice phase and the right panel shows CNO effects during the FI Training phase. Ribbons represent ±SEM.

## 4. Discussion

The present study used DREADDs-mediated chemogenetic inhibition to inactivate the PL and found that GiDREADD rats were more impulsive compared to GFP-labeled control rats following CNO administration. This effect was observed following inhibition of PL during FI training and when rats were completing the impulsive choice task. These results support the hypothesis that the PL is critical to impulsive choice *performance* and the *learning* processes by which FI training reduces impulsive choices. Reaction times at the 10s SS delay did not show any group differences, suggesting that these effects were not attributable to motoric or motivational effects of PL inhibition or CNO.

The finding that disrupting PL function increased impulsive choice during the choice procedure is consistent with some prior studies (e.g., Churchwell et al., 2009; Yates et al., 2014), however, is somewhat inconsistent with Wenzel et al. (2023) when DREADDs failed to increase impulsive choice when broadly applied to inhibit the PL (vs. just the PL-NAC circuit). Procedural differences may explain this discrepancy. Wenzel used extensive pretraining (∼180 total sessions), whereas the present study included no pre-assessments and a shorter testing period. Similarly, in set-shifting tasks, PL inhibition during limited training is disruptive, but not after extensive training (Rich & Shapiro, 2007). Thus, prolonged pretraining may reduce sensitivity to PL inhibition.

The novel finding in the present study was that PL inhibition via CNO during FI training reduced LL choices during the subsequent impulsive choice task that was administered under vehicle exposure. This outcome supports the hypothesis that learning during FI training was impaired by PL inhibition. However, this interpretation requires some caution, as the study did not assess baseline impulsive choice prior to FI training or include a control group opposite the FI training (e.g., no-delay training). Nonetheless, the finding that PL inhibition during training, when CNO was active, led to increased impulsive choice during a later test (under vehicle exposure) suggests an indirect effect on decision-making via impaired learning. This pattern is consistent with prior research showing that PL inactivation disrupts acquisition across various tasks (Balleine & Dickinson, 1998; Capuzzo & Floresco, 2020; Delatour & Gisquet-Verrier, 1996). However, for the present study, the effect involved impaired transfer of learning from the training phase (under PL inhibition) to the test phase (in the absence of inhibition). That is, the self-control promoting effects of the FI training failed to transfer to the choice task. Similar effects have been observed in contextual learning paradigms where PL disruptions during training impaired later task switching (Ragozzino et al., 2003) and condition inhibition (Meyer & Bucci, 2014).

In comparing the effects of PL inhibition across the two phases, PL inhibition during the FI training altered the slope of the choice function by increasing impulsive choices at the shorter FI delays but did not affect choice at the 30-s delay where the two choices differed only in magnitude. This suggests a more selective effect of PL inhibition during FI training on the discounting rate, where delay sensitivity was preserved despite a shift in SS choices at the shorter delays. In comparison, PL inhibition during the choice task reduced LL choices across the whole choice function. This could reflect an overall devaluation of the larger-later choice regardless of the relative value of the SS and LL options. Alternatively, the effect of PL inhibition in the choice task may have shifted preference for the SS option (like PL inhibition in the FI training task) and then subsequently retarded the ability to adapt to the new SS delays.This interpretation aligns with evidence implicating PL in goal-directed behavior, such as in studies where PL lesions impair adaptation to contingency degradation (Balleine & Dickinson, 1998) or disrupt performance on delayed spatial alternation tasks (Delatour & Gisquet-Verrier, 1996).

Our primary hypothesis posits that the timing information acquired during FI training failed to transfer to the choice procedure (Smith et al., 2019). Although the present study did not directly assess timing (e.g., via a peak interval procedure), response rates in the FI schedule provide indirect evidence of timing processes. The negatively accelerated response rates in the choice task for the group where the PL was inhibited during FI training suggest that the rats anticipated pellet delivery earlier than the programmed interval. This may reflect timing inaccuracy, where the LL delay was perceived as longer than the objective interval, which would promote impulsive choices. Another possibility is that PL inhibition during the FI training disrupted learning the value of the LL option. The FI training exposes rats to the SS and LL contingencies which may promote the integration of delay and magnitude information to guide future performance.Inhibition of the PL may prevent this encoding, leading to the lack of an increase in preference for the LL choice at the 10-s delay while remaining sensitive to changes in the SS delay during performance.

The presented study indicated a role for the PL region in impulsive choice performance and in FI Training which reduces impulsive choices. Further research will be necessary to pinpoint specific PL projections implicated. The interval timing hypothesis would posit a role of timing circuits, specifically the PL to DMS identified in Emmons et al. (2019) as containing direct PL afferents which mirror ramping activity associated with timing intervals. This circuit is also implicated in learning new goal-directed actions (Hart et al., 2018), contextual learning (Baker & Ragozzino, 2014), and value-based decision making (Choi et al., 2023), making it a prime target for refined exploration of circuits involved in impulsive choice. A baseline (pre-FI training) assessment of impulsive choice would help determine whether Gi DREADD-mediated inhibition during FI training attenuates the expected post-training reduction in impulsive choice. Future research will address these limitations with circuit-level DREADD manipulations and additional behavioral controls.

Preclinical models of behavior serve a crucial role in guiding our collective understanding of human decision-making. These studies link the granular, substrate-level mechanisms of the nervous system to the pathologies of human behavior observed in the clinic. Rodent models of impulsive choice share a high degree of translational correspondence with human decision-making. Frontal cortical regions are consistently implicated in impulsive decision-making across both human and rodent populations (Kim & Lee, 2011). In rodents, the PL is often considered an analog of the human medial prefrontal cortex (mPFC), which plays a critical role in value-based decision-making (Balleine & O’Doherty, 2010), including delayed reward value (Sackett et al., 2019). In humans, neuroimaging studies have shown that activity in the mPFC, measured via blood-oxygen-level dependent (BOLD) signals, tracks the subjective value of delayed rewards (Kable & Glimcher, 2007). Moreover, reduced BOLD responses in the mPFC have been associated with greater impulsive choice (Schueller et al., 2019). Taken together, these findings suggest a potential causal role for the PL in rats, and the mPFC homolog in humans in regulating impulsive decision-making.

Collectively, the findings of the present study demonstrate a role of the PL in the learning and performance of delay-based intervention which improve self-control. PL inhibition during the FI training or during the impulsive choice task reduced self-control, effects not explained by motivational or motoric alteration. Rats inhibited during choice performance had a reduced sensitivity to changes in the SS choice delay, suggestive of the PL’s role in cognitive flexibility. Rats inhibited during the FI training phase exhibited a steeper discounting slope relative to controls, indicating a preserved sensitivity to contingency changes. Additionally, the negatively accelerated response rates of the group which received PL inhibition during the FI intervention suggest imprecise interval timing relative to controls, consistent with the timing hypothesis. Future studies will look to dissect specific PL circuits which drive this effect on FI training to reduce impulsive choice, fostering a deeper understanding of delay discounting and its neural underpinnings.

## Notes

We have no conflicts of interest to disclose.

### Competing Interest Statement

The authors have declared no competing interest.

### Funding

- National Institutes of Health

